# Acetate promotes nutritional adaptation in *Escherichia coli*

**DOI:** 10.64898/2026.05.05.722864

**Authors:** Lucas Devlin, Virginia Oudard, Manon Barthe, Thomas Gosselin-Monplaisir, Jean-Baptiste Dupin, Florence Condamine, Jean Baudry, Muriel Cocaign-Bousquet, Pierre Millard, Brice Enjalbert

## Abstract

*E. coli* often releases acetate in its environment where it can accumulate to concentrations high enough to inhibit the bacterium’s own growth. Here, we demonstrate that acetate promotes *E. coli* adaptation to nutrient changes by accelerating growth resumption, with as little as 250 µM acetate being sufficient to shorten the lag phase by several hours. Acetate was found to be consumed via acetyl-CoA synthetase very early after the nutrient change. Transcriptomics, metabolomics and C-isotope labeling experiments show that acetate replenishes metabolic pools in the tricarboxylic acid cycle and upper glycolysis. Single-cell analyses reveal that acetate increases the adaptation speed of individual cells switching to the new nutrient. We conclude that the reuse of excreted acetate by *E. coli* facilitates nutritional adaptation by transiently replenishing central metabolite pools. This work identifies an unexpected role of acetate in the nutritional adaptation of *E. coli*, providing new insights into the physiological relevance of overflow metabolism.

**Highlights:**

- Acetate facilitates *E. coli* adaptation from one nutrient to another.
- Less than 250 µM acetate is sufficient to halve lag times.
- This effect is observed for several nutritional transitions and *E. coli* strains.
- Growth resumption is driven by the replenishment of metabolite pools through acetate assimilation mediated by acetyl-CoA synthetase.

## Introduction

The bacterium *Escherichia coli* excretes acetate during growth on many carbon sources, especially when glycolytic flux is high, as during rapid glucose consumption (Millard *et al*., 2023). This phenomenon, known as overflow metabolism, occurs in many organisms, with other metabolic end-products including ethanol in yeast and lactate in mammalian cells (Gosselin-Monplaisir *et al*., 2025a). At first glance, acetate production appears a waste of carbon and energy, and its potential inhibitory effects raises questions as to the evolutionary benefit of this production. However, acetate can also serve as a nutrient: when the glycolytic flux is low, acetate consumption alongside other carbon sources can enhance *E. coli* growth (Millard *et al*., 2023). During rapid growth on glucose, *E. coli* commonly shows net acetate excretion, followed by net consumption after glucose exhaustion (Enjalbert *et al*., 2013). This glucose-acetate transition has become a classic model for investigating nutritional adaptation (Kotte *et al*., 2010; Kremling *et al*., 2018; Pinhal *et al*., 2019). Yet, at moderate acetate concentrations, this consumption can occur without supporting measurable growth (Enjalbert *et al*., 2015). Together, these findings highlight the complex and context-dependent relationship between *E. coli* and acetate.

Here, we explored the role of acetate in *Escherichia coli* nutritional adaptation. Microorganisms such as *E. coli* live a life of feast and famine (Koch, 1971), with short periods of nutrient abundance separated by long periods of starvation. Nutritional adaptation, defined here as the ability to alternate between different nutrients, is crucial to ensure competitiveness and therefore the survival of the species in fluctuating nutritional environments. These adjustments occur when cells resume growth after starvation, when they have exhausted a preferred nutrient and must transition to other available carbon sources (i.e., diauxic shifts), or when they are abruptly transferred (switched) from one set of nutrient conditions to another. All these situations generally lead to a transient phase without growth, called the latency phase or lag phase (Monod, 1949; Madar *et al*., 2013; Moreno-Gámez *et al*., 2020). This lag phase is classically measured at the population level but the timing of adaptation can vary substantially between individual cells (Şimşek and Kim, 2018; Basan *et al*., 2020; Moreno-Gámez *et al*., 2020; Barthe *et al*., 2020). This phase is characterized by a decrease in the concentrations of most central carbon metabolites (Enjalbert *et al*., 2013; Basan *et al*., 2020) and of intracellular ATP levels (Morin *et al*., 2017; Pu *et al*., 2019). Maintenance of intracellular metabolite pools (such as phosphoenolpyruvate) can facilitate rapid growth recovery when glucose becomes available again (Xu *et al*., 2012). These observations raise a question: could the excreted acetate be reused to facilitate adaptation to changing nutrient availability, even when its concentration does not support growth?

In this work, we first compared nutritional adaptation in two settings: (i) during diauxic transitions on a mixture of two carbon sources and (ii) upon artificial switching of cells from one carbon source to another. We found that diauxic transitions were faster and showed that this is due to the presence of acetate produced during the first growth phase. Acetate was shown to shorten the lag phase across several nutritional shifts and strains, even at very low concentrations. Analysis of mutant strains showed that this benefit required acetyl-CoA synthetase (Acs). Metabolomics and isotope tracing experiments further revealed that acetate replenished central metabolite pools during early adaptation, providing a metabolic basis for its contribution to growth resumption.

## Results

### Diauxic transitions occur faster than growth recovery after artificial switching

Two experimental models are classically used to study nutritional adaptation, defined here as the capacity to switch between two carbon sources: (i) diauxic transition, where cells are grown in the presence of both nutrients and change sources when the preferred one is exhausted, and (ii) and artificial switching, where exponentially growing cells are artificially moved from one nutrient medium to the other. We evaluated adaptive efficiency under both conditions as the duration of the latency phases (lag times). The two carbon sources were glucose and xylose (Barthe *et al*., 2020). For wild type *E. coli* BW25113, the lag times for nutrient switches (0.91 ± 0.16 h) were significantly longer than for diauxic transitions (0.67 ± 0.02 h) (Figure 1A). This difference in lag times remained of the same order (on average, +240%) over a range of lag times generated by employing plasmids with varying copy numbers of the *xylA* promoter leading to the titration of the xylose-specific transcriptional factor XylR (Barthe *et al*., 2020) (Figure 1A). The fact that no lag was observed when *E. coli* populations were switched from glucose to glucose (Supplementary Figure S1) rules out cellular stress as an explanation for the difference in lag times. Meanwhile, growing *E. coli* on a mixture of glucose and xylose before switching the glucose-consuming cells to xylose (Figure 1A) only led to a slight, non-significant decrease in the lag time discrepancy (−11 ± 11%). Transcriptomic analyses (Figure 1B) confirmed that the presence of xylose during growth on glucose contributes little if at all to the lag time discrepancy (Figure 1B) : cells exponentially grown on xylose (condition X) had 100-fold higher expression of xylose-related genes (*xylA*, *xylB*, *xylE*, *xylF*, *xylG*, *xylH*, etc.) than those grown on glucose (condition G), while cells exponentially grown on glucose in the presence of xylose (condition M) showed just a twofold, non-significant (*p* > 0.05) increase in expression of *xylA*, *xylH*, and *xylB* (Supplementary Figure S2). This weak effect of xylose was further confirmed by cytometric analyses of cells grown on glucose in the presence of xylose (supplementary figure S3). Together, these results indicate that lag times for diauxic transitions are consistently shorter than those for nutrient switching, and that this difference is not attributable to the presence of the second nutrient during the first growth phase.

**Figure 1:**
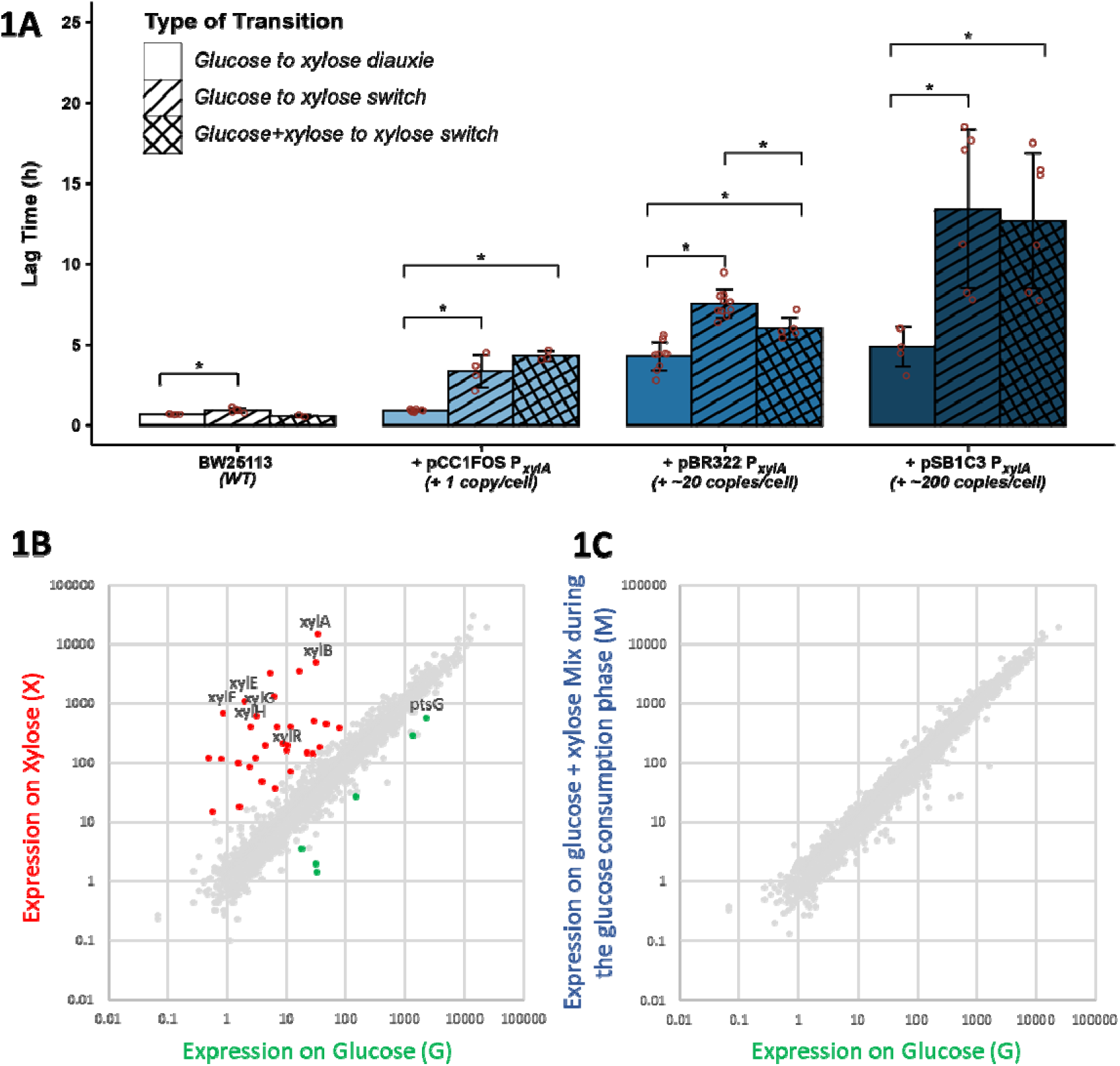
adaptation from glucose to xylose is faster following diauxic rather than artificial switching. **1A:** Lag times for E. coli strains with different levels of adaptation impairment. Growth resumption on xylose was impaired by titrating XylR with plasmids containing the XylR fixation sites of the xylA promoter (Barthe et al., 2020). BW25113 is the wild type strain without titration. Titration was achieved with plasmids pCC1FOS (single copy), pBR322 (about 20 copies) and pSB1C3 (about 200 copies) all carrying the same xylA promoter. The first column of each series represents the lag time observed during diauxic transition on a mixture of glucose and xylose, the second represents the lag time for artificial switching from exponential phase growth on glucose to xylose, and the third column represents the lag time observed when switching from exponential growth on glucose in the presence of xylose to xylose alone. Each data point is the result of five or more replicates. Error bars represent standard deviations (SD). *: significantly different values (p-value < 0.05). **1B**: Scatter plot of transcriptomic data for strain BW25113 with the titration plasmid pBR322 P_xylA_ growing exponentially on M9 medium supplemented with xylose (X) versus glucose (G) (left panel), or on glucose + xylose (M) versus glucose alone (G) (right panel). Significantly differentially expressed genes are shown in red or green, with some selected genes labelled.

### Individual cells initiate adaptation at a similar time but accumulate faster after a diauxic transition than after an artificial switch

We used the pBR322 titration plasmid expressing a fluorescent reporter under the control of the *xylA* promoter as a proxy to identify when individual xylose-adapted cells appear after artificial switching and diauxie (Figure 2). The appearance of the first adapted cells and the proportions of unadapted and adapted cells (i.e., cells expressing fluorescence under the control of the *xylA* promoter) were quantified by cytometric analysis. Expression of the reporter of xylose uptake occurred at the same time under both conditions (Figure 2B); however, the proportion of adapted cells grew faster in the diauxic condition than after switching. These results support the hypothesis that diauxic adaptation is promoted by a compound other than xylose, which is present before glucose exhaustion and whose depletion during the washing step before artificial switching reduces the cells’ ability to transition to xylose utilization.

**Figure 2:**
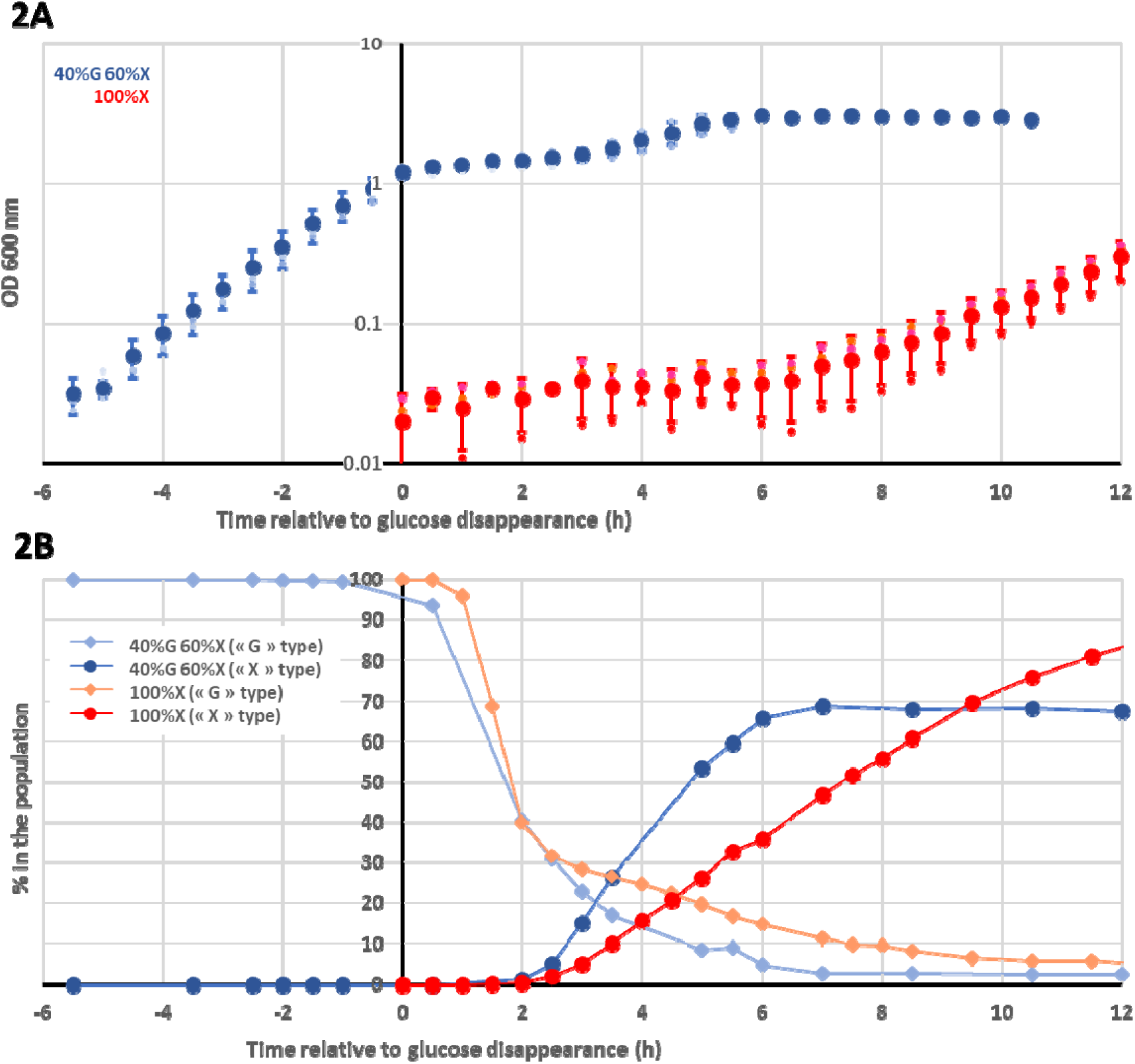
individual E. coli cells adapted to xylose appear at a similar time but accumulate faster after a diauxic transition than after an artificial switch. **2A.** Growth of E. coli strain BW25113 carrying the pBR322-based titration plasmid under glucose-xylose diauxic conditions (dark blue) or after switching from glucose to xylose (red). Zero time was defined as glucose depletion for diauxie and the start of the artificial switch, respectively (n = 3 for both). Error bars represent standard deviations (SD). **2B**. Cell cytometric profiles under the same conditions. The xylA promoter carried by plasmid pBR322 controls the expression of the red fluorescent reporter mRFP1. Fluorescent cells (i.e., xylose-metabolizing cells) were defined by their profiles on xylose alone and presented as “X” cells. They are represented by dark-blue circles during the diauxic transition and red circles during the artificial switch. Non-fluorescent cells were defined based on their profiles on glucose alone and presented as “G” cells. They are represented by light-blue diamonds during the diauxic transition and salmon-pink diamonds during the artificial switch. The sum of the G and X cell types does not add up to 100% because cells with intermediate profiles, typically corresponding to cells transitioning from G to X profiles, are not quantified here.

### Acetate promotes nutritional adaptation

Based on the previous observations, we hypothesized the adaptation-enhancing compound might be acetate. Indeed, acetate is produced during growth on glucose (Enjalbert et al., 2015) and it is therefore present during diauxic growth, but lost during the washing step when the cells are switched onto fresh medium. We therefore tested whether addition of acetate to the new xylose medium reduced the lag times after artificial switching. At acetate concentrations equivalent to those under diauxic conditions (4 mM; Enjalbert *et al*., 2015), the lag time was reduced nearly three-fold compared with control (acetate-free) conditions (Figure 3A, left panel, condition 4 mM). Surprisingly, the same significant reduction in lag time was observed at acetate concentrations as low as 250 µM, while increasing the acetate concentration did not further improve nutritional adaptation. We then assessed the generality of this finding using two other nutritional transitions (galactose to fucose and N-acetylglucosamine (NAG) to malate) that showed sufficiently long lag phases in the wild-type BW25113 strain to measure a putative reduction in lag induced by acetate. A similar effect of acetate with a similar threshold concentration of 500 µM was observed in both transitions (Figure 3A, mid and right panels). This very low acetate concentration is insufficient to support measurable growth (Enjalbert *et al*., 2015). A similar effect of acetate was observed in both microplate and flask experiments (supplementary figure S4), albeit with a shorter lag phase in microplates than in flasks, as previously described (Barthe *et al*., 2020). To investigate whether this benefit extended across *E. coli* strains, we tested BW25113, M1/5, MG1655, and Nissle 1917 using a glucose-to-fumarate transition. This transition was selected because all four strains could grow on both substrates and exhibited sufficiently long lag phases to assess the effect of acetate. Acetate shortened lag times in all four strains (Figure 3B).

**Figure 3:**
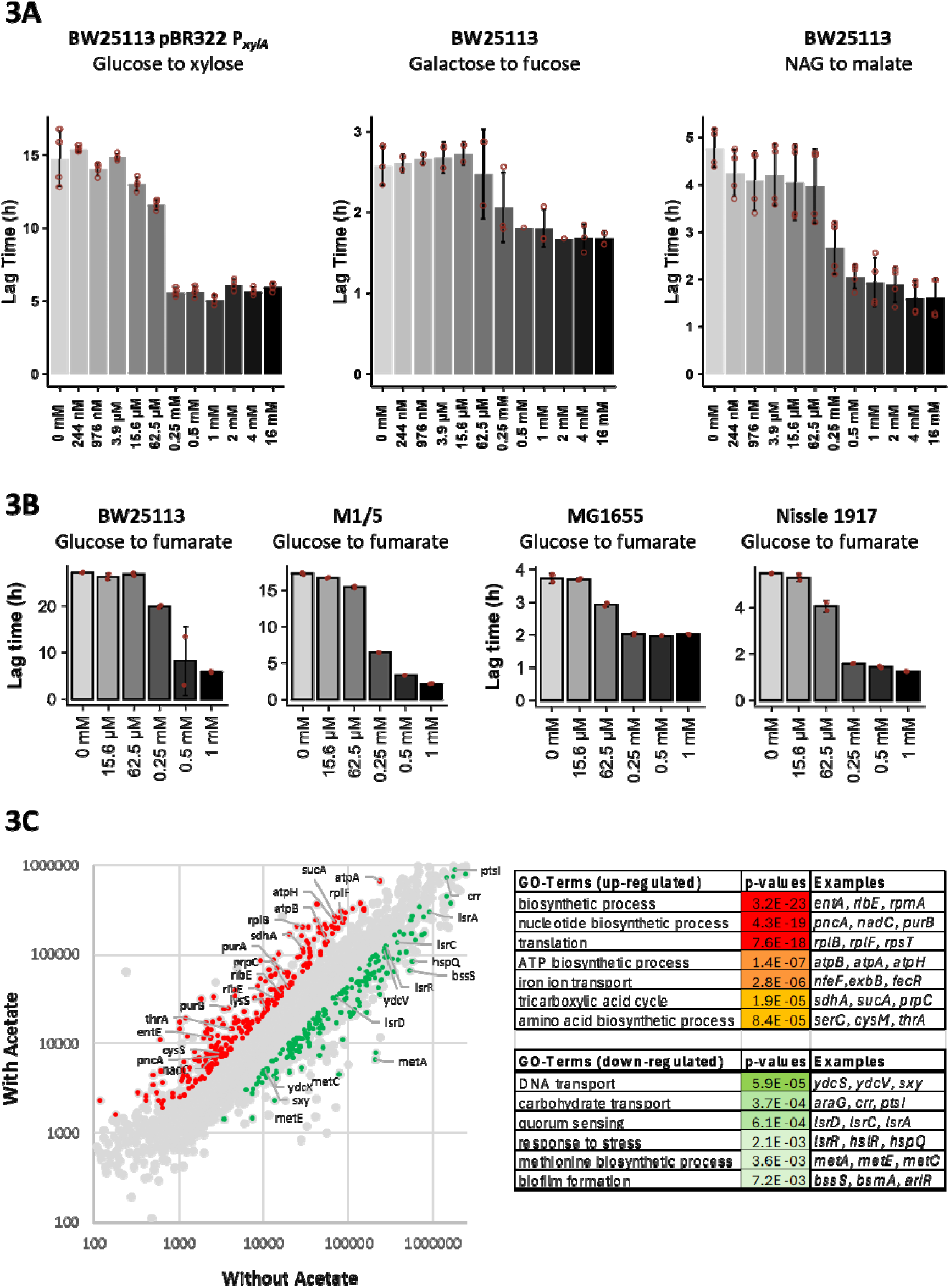
acetate promotes nutritional adaptation. **3A**: lag times were measured in E. coli BW25113 cultures switched from one substrate to another in the presence of 0 to 16 mM acetate: left panel, results for BW25113 carrying the titration plasmid pBR322 P_xylA_ switched from glucose to xylose; middle panel, wild type BW25113 switched from galactose to fucose; right panel, wild type BW25113 switched from N-acetylglucosamine to malate. N = 2 to 4 repetitions are shown as open red circles for each condition. Error bars represent standard deviations (SD). **3B**: The same experiment was performed using glucose-to-fumarate switches in strains BW25113, M1/5, MG1655, and Nissle 1917. Error bars represent standard deviations (N=2). **3C**: transcriptomic analysis of wild type BW25113 30 min after switching from N-acetylglucosamine to malate in the presence or absence of 3 mM acetate. Left panel: scatterplot of gene expressions with significantly differentially expressed genes in red (3-fold higher expression in the presence of acetate) or green (3-fold lower expression in the presence of acetate), and with some selected genes labelled. Right panel: gene ontology analysis of the red-and green-highlighted genes.

Finally, transcriptomic analysis comparing cells 30 min after switching from NAG to malate was performed with or without acetate (Figure 3C). Since 0.5 mM acetate was sufficient to achieve maximal lag reduction at an OD600 of 0.1, and since a six-fold higher cell concentration (OD600 = 0.6) was required for the transcriptomic analysis, we chose an acetate concentration of 3 mM to ensure the same acetate availability per cell. This revealed a strong up-regulation in the presence of acetate for genes implicated in translation, nucleotide biosynthesis and TCA. These results all indicate that acetate boosts nutritional adaptation in distinct *E. coli* strains, even at very low concentrations – far lower than those required to sustain growth.

### Acetate accelerates individual cells adaptation to the new nutrient

To explore how acetate promotes nutritional adaptation in cells, we first investigated individual cells adaptation to a new substrate. Flow cytometry experiments were performed after switching *E. coli* strain BW25113 containing the reporter gene mNeonGreen under control of the *fucA* promoter (Figure 4A) from galactose to fucose. The proportion of fluorescent cells expressing the reporter (i.e. metabolizing fucose) increased more rapidly in the presence of 0.5 mM acetate (Figure 4B). Comparing the proportions of cells derived from non-fluorescent versus already adapted cells (Figure 4C and Supplementary Figure S5 for the replicates) shows that adaptation in the presence of acetate occurred in the first hour after switching, and one hour later without acetate. Interestingly, a similar plateau was observed 2 h after the switch with or without acetate. Six hours after the switch, about 40% of the initial population remained unadapted in both cases. This confirms the previous findings (Figure 1C) that acetate accelerates adaptation of *E. coli* to the new nutrient, but does not increase the overall fraction of cells able to adapt.

**Figure 4:**
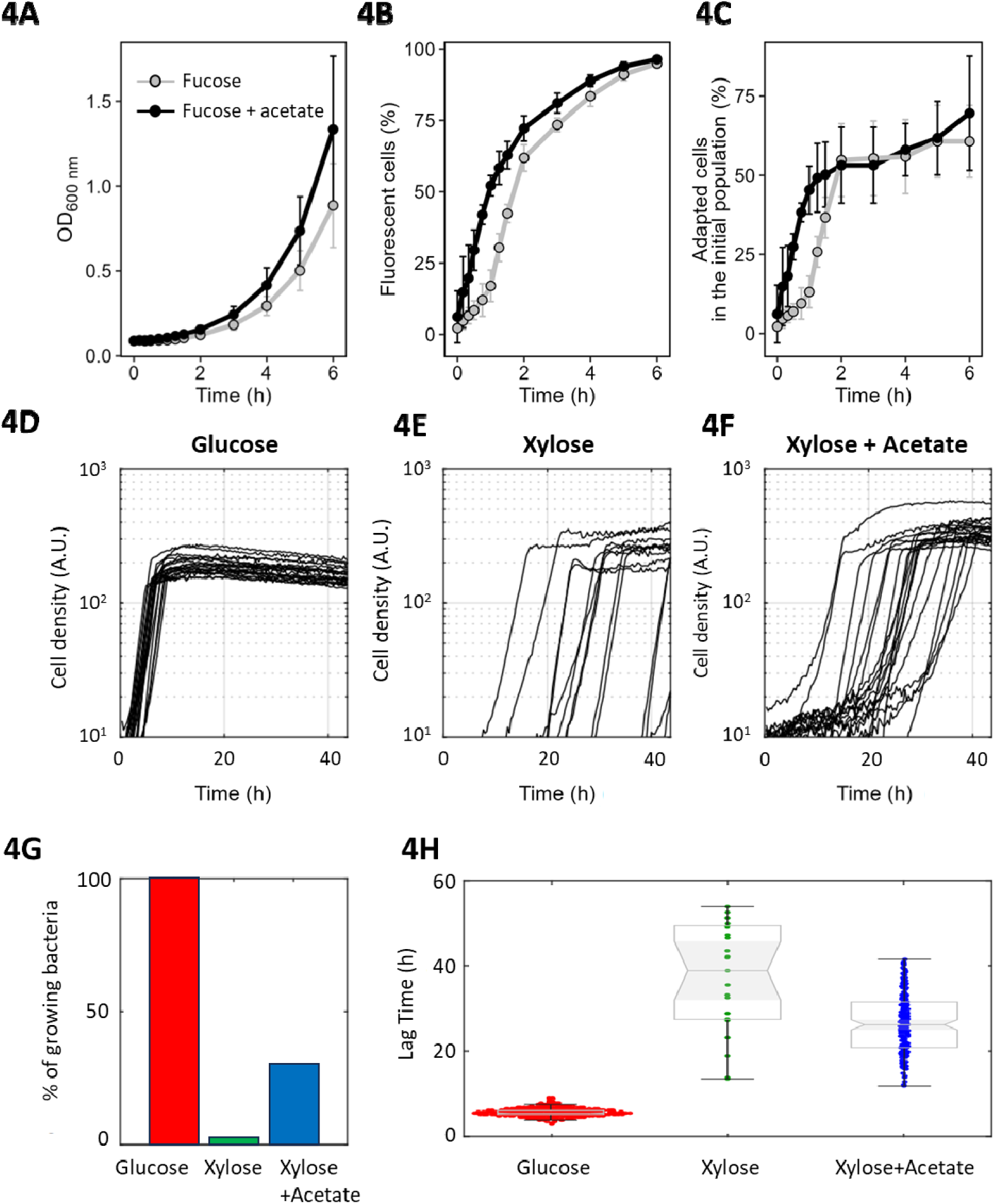
acetate accelerates the adaptation of individual cells. Strain BW25113 pCC1FOS P_fucA_-mNeonGreen P_j23119_-mScarlet-I was switched from galactose to fucose with or without 0.5 mM acetate (black and grey respectively) (n = 3; error bars represent standard deviations). Time-evolutions of total biomass (**4A**), fluorescent subpopulation (**4B**) and the proportion of adapted cells relative to the initial population (**4C**). (4D–4H) Droplet experiments with strain BW25113 carrying the titration plasmid pBR322 P_xylA_. Bacteria were precultured in glucose and individual cells were switched to droplets containing either glucose (**4D**), xylose (**4E**), or xylose + 0.5 mM acetate (**4F**). The growth curves of individual droplets were measured by monitoring cell autofluorescence (cell density expressed in arbitrary units; 20 randomly selected curves shown per condition). (**4G**) Percentage of bacteria inoculated into droplets that initiated divisions within 40 h (n = 3). (4H) Boxplot comparisons of the lag time under the studied conditions (n = 440 for glucose, n = 25 for xylose, and n = 199 for xylose + acetate).

To investigate the timing of individual cell adaptations to xylose, cells expressing mNeonGreen under the *xylA* promoter were individually switched from glucose to 50 µl droplets containing either glucose (Figure 4D), xylose (Figure 4E) or xylose + 0.5 mM acetate (Figure 4F). This allowed us to track when each cell resumed growth. While cells switched from glucose to glucose all resumed growing instantaneously, xylose-switched cells rarely adapted (Figure 4G), with highly variable timing of growth resumption for those that did (Figure 4E). As observed for the galactose-fucose switch, acetate facilitated adaptation from glucose to xylose, reducing the average lag time, and reducing lag time variability (Figure 4H). However, acetate did not significatively affect the subsequent growth rate (0.51 ± 0.07 h^−1^ with acetate versus 0.45 ± 0.10 h^−1^ without). These results indicate therefore that acetate accelerates adaptation of individual cells to the next nutrient, thereby reducing the overall lag at the population level.

### Acetate consumption during nutritional adaptation feeds the TCA cycle and replenishes tricarboxylic acid pools

Since acetate accelerates adaptation to new nutrients, we hypothesized that it is consumed following depletion of the first nutrient and investigated the timing of its consumption. We measured the evolution of extracellular metabolite concentrations by NMR after switching from NAG to malate, which induces a long lag in the wild type strain, facilitating the experiments (Figure 5A-C). To match the conditions used for the subsequent metabolomic analysis, the inoculation on malate was performed at OD_600nm_ = 0.6. The corresponding experiment at OD_600nm_ = 0.1 is provided as Supplementary Figure S6. In both cases, added acetate at 0.5 mM was consumed in less than 60 min after switching and reduced the lag time (Figure 5A & 5B; Supplementary Figure S6). At 3 mM, acetate was also rapidly consumed, followed by acetate production before its subsequent exhaustion once the population had resumed growth on malate (Figure 5C).

**Figure 5:**
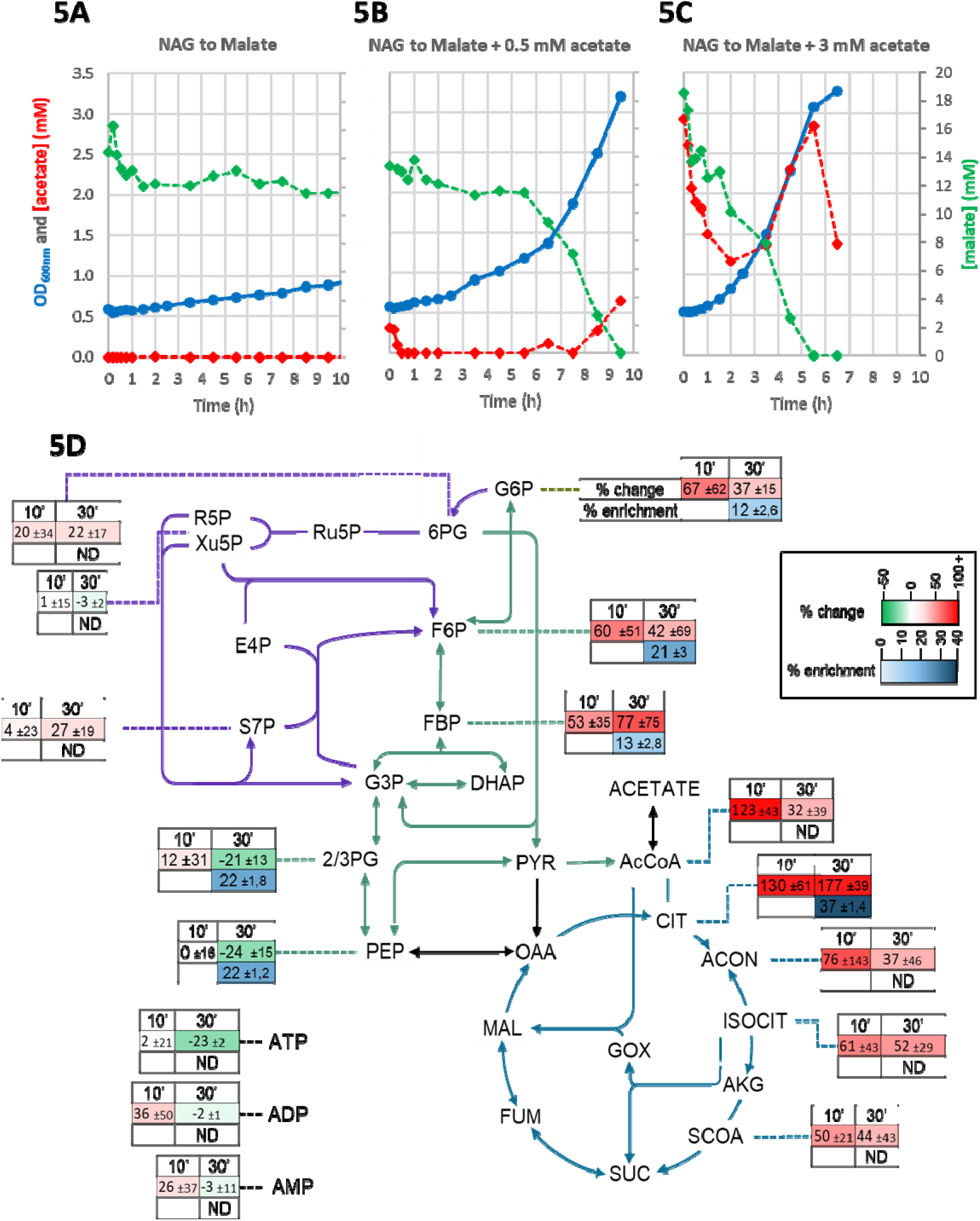
acetate consumption and utilization during switching from NAG to malate. **(5A–5C)** Extracellular metabolites during switching from N-acetylglucosamine to malate in flasks. Biomass (optical density; blue dots), acetate concentration (red dots) and malate concentration (green dots) as a function of time, in the absence of acetate (**5A**), with 0.5 mM acetate added to the malate (**5B**) and with 3 mM acetate added to the malate (**5C**; note that growth ended earlier under this condition because growth resumed earlier, while 10 hours were insufficient for growth resuming in the absence of acetate). (**5D**) Comparison of metabolic pools in the central carbon metabolism in the presence or absence of acetate. Differences are shown as percentages 10 and 30 min after switching from N-acetylglucosamine to malate for the wild-type strain. Percentage enrichment in C in the presence of 3 mM C-labeled acetate is shown as white to dark-blue shading. ND: not quantifiable. These are the results of three independent experiments.

Metabolomic analysis showed that acetate was found to significantly increase glycolytic, acetyl-CoA, and tricarboxylic acid pools (citrate, aconitate, isocitrate, succinyl-CoA) within the first 10 min of switching from NAG to malate (Figure 5D). These results suggest an increase in TCA fluxes and thus of ATP production. However, acetate did not significantly change ATP concentrations and adenylate energy charge (Figure 5D and Supplementary Figure S7), suggesting that the newly produced ATP is readily consumed in other cellular processes.

To confirm that pool replenishing is directly related to acetate consumption, we measured the mean ^13^C-enrichment of central metabolites 30 min after the switch in ^13^C-isotope labeling experiments with 3 mM ^13^C-labeled acetate (Figure 5D; blue values). ^13^C-enrichments of citrate reached 37 ± 1%, demonstrating that acetate feeds the TCA cycle, in keeping with the metabolomics data. Interestingly, ^13^C-enrichments of glycolytic intermediates of between 12 and 22% were also observed, indicating that acetate also feeds the upper part of central metabolism through neoglucogenesis. Acetate therefore acts by increasing the acetyl-CoA pool within the first minutes of nutrient switching, and this then propagates through the central metabolism to provide energy and replenish other central metabolic pools. Finally, when acetate was replaced by other TCA cycle metabolites (fumarate, malate or succinate) in the same galactose-to-fucose transition as in Figure 3A, a similar reduction in lag was observed (Supplementary Figure S8). This further supports the idea that TCA cycle replenishment, rather than a specific property of acetate, is important for the observed faster nutritional adaptation.

### The boosting effect of acetate is dependent on acetyl-coA synthetase

Previous studies highlighting the essential role of the Pta-AckA pathway (Figure 6) in acetate production and consumption during growth on glycolytic nutrients (Enjalbert *et al*., 2017; Millard *et al*., 2023) raise questions about the function of the alternative acetyl-coA synthetase (Acs) pathway. NAG to malate switching experiments were repeated with different mutants and the booster effect of acetate disappeared only in the Δ*acs* strain (Figure 6). A transcriptomic analysis revealed no significantly enriched cellular functions in the Δ*acs* mutant in response to acetate, suggesting that acetate promotes nutritional adaptation primarily through its assimilation and metabolism, with no evidence of an additional metabolism-independent effect (Supplementary Figure S9). Acs has a higher affinity for acetate than does the Pta-AckA pathway (Kumari *et al*., 2000; Valgepea *et al*., 2010) and has been previously described to be induced after glucose depletion (Enjalbert *et al*., 2013; Enjalbert *et al*., 2015). Our results therefore indicate that the early consumption of small amounts of acetate that accelerate nutritional adaptation requires the Acs pathway.

**Figure 6:**
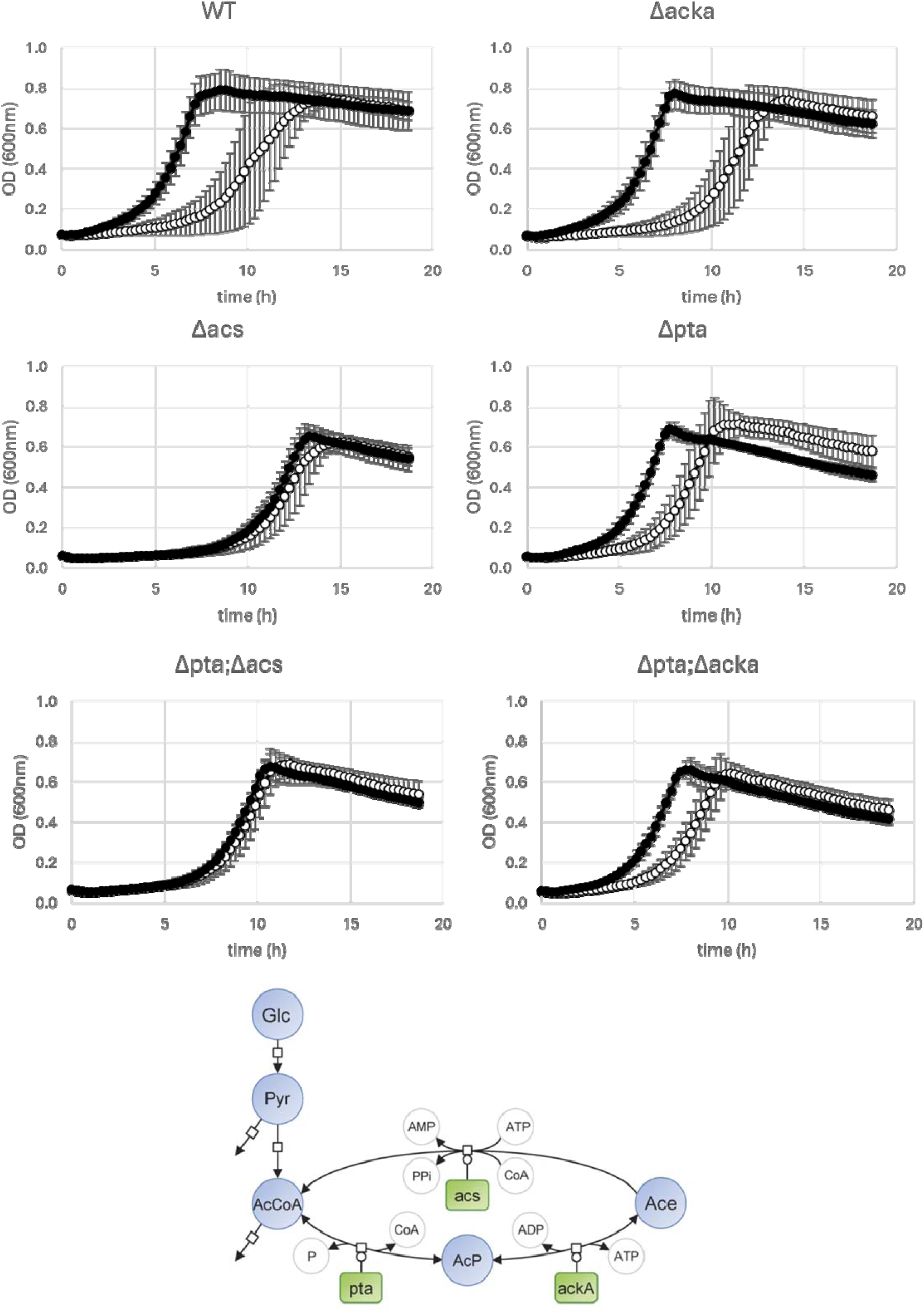
acs is required for acetate boosting of nutritional adaptation. Comparisons of the time evolution of biomass after a switch from N-acetylglucosamine to malate in the presence (black circles) or absence of 0.5 mM acetate (open circles) for wild-type E. coli BW25113 and for various single or double mutants of acetate metabolic pathways (shown in the lower panel). Error bars represent standard deviations (N=3).

## Discussion

The results of this study reveal how acetate plays an unexpected and crucial role in boosting *E. coli* nutritional adaptation: very small amounts of acetate, insufficient to sustain growth, promote adaptation to the second nutrient in various nutrient switches. This finding extended across different nutritional transitions and *E. coli* strains. The positive effect of acetate stems from its early utilization following immediate expression of Acs to promote its catabolic utilization through the TCA cycle. This replenishes central metabolic pools and provides carbon and energy, promoting the rewiring of the cell machinery to the next nutrient, and thus reducing lag times. This also explains why the lags observed in glucose-related diauxic transitions are shorter than those for nutrient switches, because acetate is already present in the former situation when cells have to reorganize their metabolism in preparation for the second nutrient.

Our results support a mechanism involving acetate utilization and the replenishment of central metabolite pools. The requirement for Acs, together with the incorporation of acetate-derived carbon into central metabolites, links the acetate adaptation-promoting effect to its utilization. Consistent with this interpretation, transcriptomic analysis detected no significant enrichment of biological processes among acetate-responsive genes in the Δ*acs* strain. The ability of other TCA cycle metabolites to shorten lag times further supports a contribution of central metabolism to growth resumption. Because acetate accumulates during growth on carbon sources such as glucose, its subsequent reuse can provide a readily available resource during nutritional transitions. The plateau in lag reduction is consistent with acetate alleviating a transient metabolic limitation, after which other processes constrain growth resumption. Our findings thus identify a physiological benefit of acetate reuse: facilitating adaptation to alternative carbon sources.

Previous studies have linked lag duration to the metabolic state before the nutrient shift, including carbon influx and the balance between respiratory and fermentative metabolism (Basan *et al*., 2020; Fuentes *et al*., 2021). Our findings identify extracellular acetate availability as an additional factor influencing adaptation. Madar *et al*. (Madar *et al*., 2013) observed that genes involved in utilization of the new carbon source are expressed before measurable biomass growth, followed by activation of a broader growth program. Acetate consumption may therefore shorten this preparatory phase by supplying precursors and energy for the synthesis of enzymes required to utilize the new nutrient.

Metabolomics revealed acetate increases central metabolite pools after the nutritional switch, while ^13^C-labeling experiments confirmed that acetate-derived carbon entered TCA cycle and gluconeogenic intermediates. These findings extend our previous observations of acetate utilization after glucose exhaustion (Enjalbert et al., 2017). The increased expression of TCA cycle genes, together with the incorporation of acetate-derived carbon into this pathway, This indicates an energetic contribution of acetate during adaptation, through the generation of reducing equivalents that support respiratory ATP synthesis. This is further supported by the ability of other TCA cycle metabolites to shorten lag times. The absence of a detectable increase in ATP concentration or adenylate energy charge does not contradict this interpretation: ATP generated from acetate metabolism may be consumed during adaptation without accumulating. Such coordination between ATP production and consumption is consistent with the maintenance of adenylate energy charge under fluctuating substrate availability previously reported in *E. coli* (Vasilakou *et al*., 2020).

Acetate has long been recognized for its inhibitory effects on growth (Luli, 1990; Pinhal *et al*., 2019). We previously demonstrated that acetate can enhance growth at low glycolytic flux (Millard *et al*., 2023). Here, we identify an additional beneficial role in nutritional adaptation. One of our most surprising observations in this study was that even trace amounts of acetate can have a dramatic effect on metabolic lags, emphasizing the complex pattern of interactions between acetate and *E. coli*. These findings suggest that excreted acetate can act as an extracellular carbon reserve for *E. coli*, available for reuse during nutritional transitions to replenish central metabolite pools and facilitate growth resumption. In keeping with previous studies (Şimşek and Kim, 2018; Dal Co *et al*., 2019), we observed substantial variability in the timing of individual cells’ nutritional adaptation. Acetate reduced this variability and shortened the overall lag time, without increasing the fraction of cells in the initial population capable of adapting.

Our findings are important both for basic research and for applications. Our study highlights a very easy way to substantially reduce lag times, which should increase the profitability and sustainability of bioprocesses. Fundamentally moreover, this work alters our perception of the relationship between *E. coli* and acetate. In fluctuating environments such as the gut, this relationship could be crucial for *E. coli* survival. The role of acetate in complex microbiomes therefore warrants further investigation. Another avenue of interest is the broader role of the exometabolome in growth resumption. While acetate appears to be a prominent contributor, other compounds present in the extracellular medium may also contribute to growth resumption. Finally, since similar overflow mechanisms exist in yeast (with ethanol) and mammalian cells (with lactate) (Gosselin-Monplaisir *et al*., 2025a), it would be interesting to investigate whether the same boosting effects are observed for the overflow products in these organisms.

## Acknowledgements

This work was supported by the ANR project JANUS (ANR-19-CE43-0004-01). The authors thank Toulouse White Biotechnology and its staff for providing access to their cytometry facilities. LD was supported by the MICA department of INRAE. PMI and TGM were supported by both the MICA department of INRAE and Région Occitanie (grant COCA-COLI). PMI was also supported by the MITI program of CNRS (grant METABO-FLOW). The authors thank Lindsay Peyriga and MetaboHub-MetaToul (Metabolomics and fluxomics facilities, Toulouse, France, https://mth-metatoul.com), part of the French National Infrastructure for Metabolomics and Fluxomics (http://www.metabohub.fr) funded by the ANR (MetaboHUB-ANR-11-INBS-0010), for access to NMR facilities. The authors are grateful to the following INSA Toulouse students and the Department of Biological Engineering for help with the microrarray experiments: Eva Antico, Tristan Baeumlin, Camille Dessemond, Esteban Duneau, Margot Duhamel, Hermeline Frenoy, Florian Jabally, Emma Landolt, Lorenzo Molinengo, Océane Renard, Zawata-Afnan Sharara and Julie Souloumiac.

## Methods

### Strains and plasmids

*Escherichia coli* K-12 BW25113 *rrnB3* Δ*lacZ4787 hsdR514*Δ*(araBAD)567* Δ*(rhaBAD)568 rph-1* was chosen as the wild-type model strain. Combinations of Δ*pta*, Δ*ackA*, and Δ*acs* mutants were constructed from wild-type *E. coli* K-12 BW25113 from the KEIO collection (Baba *et al*., 2006) after removal of the kanamycin cassette (Datsenko and Wanner, 2000). The *pta* and *acs* genes were individually deleted by CRISPR-Cas9 genome editing using the method and plasmids described in Jiang *et al*. (Jiang *et al*., 2015). The deletions were confirmed by genome sequencing or by PCR.

Plasmid pSB1C3 P_ihfb_-mTagBFP_BBa_0015_P_xylA_-mRFP1 was obtained from Barthe *et al*. (Barthe *et al*., 2020). The other plasmids were constructed by Gibson assembly (NEBuilder® HiFi DNA Assembly from New England Biolab, France) performed following the protocol recommended by the manufacturer to combine a plasmidic chassis and double reporter system from the pSB1C3 plasmid. Chassis pBR322 was obtained from New England Biolabs (ref N3033S) but with the chloramphenicol resistance cassette from pSB1C3 instead of its original ampicillin resistance cassette. Chassis pCC1FOS was obtained from from Epicentre^TM^. mTagBFP was replaced by mScarlet-I in the pCC1FOS chassis. mRFP1 was replaced by mCherry in the pBR322 chassis and by mNeonGreen in the pCC1FOS chassis. The *ihfb* promotor was replaced by the synthetic promotor P_J23119_ from the Anderson promotor collection fused to a strong RBS derived from pDawn (Part:BBa_K3288007 by Joshua Nam). P_xylA_ was replaced by the fucose specific promoter P_fucA_, amplified from *E. coli* BW25113 genomic DNA.

### Growth media

The strains were grown in M9 minimum medium containing Na_2_HPO_4_·12H_2_O (48.5 mM), KH_2_PO_4_ (22 mM), NaCl (8.6 mM), NH_4_Cl (38.1 mM), MgSO_4_·7H_2_O (2 mM), CaCl_2_·2H_2_O (30 µM), Na_2_EDTA·2H_2_O (40.3 µM), ZnSO_4_·7H_2_O (15.6 µM), CoCl_2_·6H_2_O (1.3 µM), MnCl_2_·4H_2_O (5.1 µM), H_3_BO_3_ (16.2 µM), Na_2_MoO_4_·2H_2_O (1.7 µM), FeSO_4_·7H_2_O (10.8 µM), CuSO_4_·5H_2_O (1.2 µM), and thiamine-HCl (0.1 g/L), and carbon sources at 90 mMeqC (with 40% glucose + 60% xylose for diauxic growth). For the culture of strains containing plasmids, M9 medium was supplemented with 20 mg·L^−1^ chloramphenicol (except for cytometric experiments that were performed without chloramphenicol for the pCC1FOS containing cells).

### Flask cultures and switch

Cells were pre-grown overnight in 50 mL flasks containing 10 mL of M9 minimal medium supplemented with the initial carbon substrate (glucose, galactose, or NAG) at 37°C and 150 rpm. After overnight growth, cells were centrifuged at 4000 rcf for 2 min, the supernatant was discarded, and the pellet was resuspended in fresh M9 medium with the same initial carbon substrate. This resuspension was used to inoculate a 250 mL flask containing 50 mL of fresh M9 minimal medium supplemented with the same initial carbon substrate at OD_600 nm_ = 0.1. Cells were grown until they reached an OD_600 nm_ of 0.8–1.2. At this point, cells were centrifuged again at 4000 rcf for 2 min, the supernatant was discarded, and cells were resuspended in 10 ml pre-warmed fresh M9 minimal medium supplemented with the second carbon substrate (xylose, fucose, or malate). This resuspension was centrifuged again at 4000 rcf for 2 min, the supernatant was discarded, and cells resuspended in 50 ml pre-warmed fresh M9 minimal medium supplemented with the same carbon substrate in a new 250 mL flask containing at an initial OD_600nm_ of 0.1. Optical density at 600 nm was measured using a Jenway 7200 spectrophotometer (Jenway™, UK) to monitor growth.

For diauxic growth, cells were pre-grown overnight in M9 minimum media supplemented with glucose, then centrifuged and resuspended in fresh new media supplemented with a glucose-xylose mix. This resuspension was used to inoculate a flask containing the glucose-xylose mix at an initial OD_600nm_ of 0.1 to monitor growth.

### Microplate cultures and switch

Cells were pre-grown overnight in 50 mL flasks containing 10 mL of M9 minimal medium supplemented with the initial carbon substrate (glucose, galactose, or NAG) at 150 rpm. After overnight growth, cells were centrifuged at 4000 rcf for 2 min, the supernatant was discarded, and cells were resuspended in fresh M9 minimal medium with the same initial carbon substrate. This resuspension was used to inoculate a 250 mL flask containing 50 mL of fresh M9 minimal medium supplemented with the same initial carbon substrate, at an initial OD_600 nm_ of 0.1. Cells were grown until they reached an OD_600nm_ of 0.8–1.2. In parallel, 96-well microplates were prepared with 100 µL per well of M9 minimal medium containing a 2X concentration of the second carbon substrate (xylose, fucose, malate, or fumarate) and a 2X concentration of acetate. Outer wells were used as blanks. Cells grown to OD_600 nm_ 0.8–1.2 were centrifuged at 4000 rcf for 2 min, the supernatant was discarded, and cells were washed and resuspended in fresh M9 minimal medium without any carbon source. This cell suspension (2X concentration) was then used immediately to inoculate the microplate wells in duplicate by adding 100 µL per well, achieving a final initial OD_600 nm_ of 0.2 per well. Microplates with lids were read at OD_600nm_ at 17 min intervals for 24 h at 37°C with agitation on a Versamax microplate reader (Molecular devices, USA).

### Batch cultures

Bioreactor batch cultures were performed in a Sartorius Biostat B plus bioreactor in 1 L of M9 medium with 90 mMeqC of glucose and/or xylose mixes at 37 °C and pH 7. Non-limiting aeration conditions were obtained with an air flow of 0.35 L·min^−1^ and adaptation of stirring to maintain pO2 > 20%. Growth was assessed by OD_600 nm_ measurements at 30 min intervals with a LibraS4 spectrophotometer (Biochrom, UK).

### Droplet cultures

Cells precultured at 37°C on glucose M9 medium were rinsed twice and resuspended in M9 medium supplemented with either glucose, xylose or xylose plus 0.5 mM acetate. Three microfluidic chips are used, one for each medium. Droplets were formed in the microfluidic chip and imaged according to the protocol described in Eyer *et al*. (Eyer *et al*., 2017). Briefly, drops of around 50 pL were formed and then 2D confined between two plastic coverslips. They were then imaged using an inverted optical microscope (Nikon TiE) at 37°C with a x20 objective (Nikon, NA 0.75) every 20 min for 60 h, with an ORCA flash4.0. Focus was maintained using the Perfect Focus system (PFS, Nikon). The three chips were imaged in parallel, in bright-field and fluorescence (CFP and mCherry channels). One image corresponds to around 200 drops, and between 5 and 10 images were taken per chip. The initial inoculum was chosen to have a majority of single bacteria per drop at the start of the experiment, with an average of around 0.3 bacteria per drop. The films were analyzed using a Matlab program (https://github.com/ESPCI-LCMD/MiMB) (Bounab *et al*., 2020). Drops that did not move by more than 25% of their radius were tracked over time in bright-field images. Fluorescence images were used to characterize bacterial growth, particularly the CFP channel. Illumination inhomogeneities were corrected across all images using BaSiC (Peng *et al*., 2017) and the autofluorescence intensity was averaged for each drop. Drops without bacteria were used to calculate the background intensity, which was averaged per image and subtracted from the signal of drops containing bacteria. The lag time was defined here as the time required for the CFP auto-fluorescence signal to reach an arbitrary threshold of 70.

### Lag calculations for flask cultures

The term “lag” refers to the time required for cell populations to reach exponential phase after exhaustion or removal of the first nutrient (Enjalbert *et al*., 2015; Basan *et al*., 2020; Barthe *et al*., 2020). The formulas used for switching and diauxic lags are described in Enjalbert *et al*., 2015 and Barthe *et al*., 2020, respectively.

### Lag calculations for microplates cultures

Growth curves were analyzed in R with the *gcplyr* package (Blazanin, 2024). Growth rates were calculated using the *calc_deriv* function (with *window_width_n* set to *5*), and lag times were calculated using the *lag_time* function.

### Flow cytometry

Sampled cells were stored at 4°C. Cells were then diluted in filtered PBS to obtain a concentration of 2 × 10^6^ cells·mL^−1^. Fluorescence intensities of individual cells were measured with a Masquant VYB cytometer (Miltenyi Biotec, Germany) equipped with a 561 nm yellow laser for excitation of mRFP1 and mScarlet-I, a 405 nm purple laser for excitation of mTagBFP and a 488 nm blue laser for excitation of mNeonGreen. Data acquisition was set at 20 000 events per sample. FlowJo X software was used for cytometry data analysis. A gate was created in the dot plot of the forward scatter channel (FSC-H) versus the side scatter channel (SSC-H) to distinguish bacteria from technical noise. A second gate was created in the dot plot of the SSC-H versus the SSC-A to select single cells. mTagBFP fluorescence emissions were analyzed with the V1-H channel (BP452/45nm), mRFP1 emissions with the Y2-H channel (BP615/20 nm), mScarlet-I emission with the Y1-H channel (BP586/15 nm) and mNeonGreen emissions with the B1-H channel (BP525/50 nm).

The proportion X_t_ of adapted cells from the initial population (i.e., the proportion of cells from the inoculum that had switched and resumed growth on fucose at time t) was calculated as follows. The number of cells that adapted between t_−1_ and t, denoted S_t_, was determined as the difference between the total number of fluorescent cells (i.e., fucose-adapted cells, A_t_) and the number of fluorescent cells resulting from the division of previously adapted cells (G_t_):

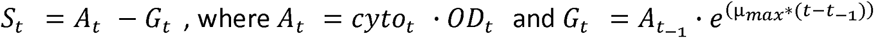

Here, *cyto_t_*represents the fraction of fluorescent cells as determined by flow cytometry, and μ_max_ is the maximal growth rate on fucose, determined from cells growing exponentially on fucose, which was estimated by linear regression of *ln(A_t_)*. The cumulative proportion of adapted cells was then calculated as:

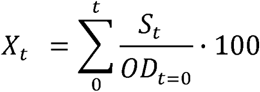

### Extracellular metabolome

Extracellular concentrations of nutrient were quantified in 180 µL of filtered broth (0.2 μm syringe filter, Sartorius, Germany) by 1D ^1^H-NMR on a Bruker Avance 500 MHz spectrometer equipped with a 5-mm z-gradient BBI probe (Bruker, Germany), as described previously (Millard *et al*., 2021).

### Intracellular metabolome

To quantify the concentrations and ^13^C-contents of intracellular metabolites, samples were collected by fast filtration as described in (Millard *et al*., 2014). For acyl-CoAs (acetyl-and succinyl-CoA), 5 mL of culture broth were harvested by vacuum filtration (0.45 μm Sartolon polyamide, Sartorius, Germany), washed with 5 mL of the same medium with reduced concentrations of phosphate and sulfate salts, and rapidly quenched in liquid nitrogen. For other central metabolites, 200 µL of culture broth were harvested by vacuum filtration (0.45 μm Sartolon polyamide, Sartorius), washed with 1 mL of the same medium with reduced concentrations of phosphate and sulfate salts, and quenched in liquid nitrogen. Intracellular metabolites were extracted by incubating filters in 5 mL of extraction solution (methanol/acetonitrile/water 40/40/20 with 0.5 % formic acid) at −20°C for 1 h. Cell extracts were centrifuged to remove cell debris, and supernatants were dried overnight under vacuum and stored at −80°C until MS analyses.

For LC-MS analyses, dried extracts were resuspended in 200 µL of milliQ water. Pools of acyl-CoAs were quantified as described in (Gosselin-Monplaisir *et al*., 2025b), and other central metabolites were analyzed as described in (Vogeleer *et al*., 2024). The raw data were processed using Skyline v24.1.0. For isotopic measurements, the mean ^13^C-enrichment was calculated from the intensities of mass fractions after correction for naturally occurring isotopes and for the isotopic purity of the ^13^C-labeled acetate (99 %), using IsoCor v2.2.2 (Millard *et al*., 2019)(https://github.com/MetaSys-LISBP/IsoCor).

### RNA-Seq transcriptome

Experiments were performed with cells obtained from either exponential phase on glucose (G), exponential phase on xylose (X) or a glucose-xylose mix during the glucose exponential phase (M) cultures. Three independent replicates were carried out. Cell samples were processed to extract RNA by TRIzol/phenol treatment (Esquerre *et al*., 2014). ERCC (ERCC RNA Spikes-In Control Mix 1, AMBION) was added before ribodepletion with 1 µL of ERCC diluted 1/10 to 5000 ng of total RNA. Ribodepletion was performed with the Pan-Karyote SiTools ribo POOL kit (siTOOLs Biotech). RNAseq analysis was performed in IonTorrent (Ion S5, Thermofischer) on the GETbiopuce platform (Nguyen *et al*., 2022). Data were normalized using the R package, according to the TMM_exact methodology (Roux *et al*., 2025). The transcriptomics data can be downloaded from the ArrayExpress database (www.ebi.ac.uk/biostudies/arrayexpress) under accession number E-MTAB-14542.

### Microarray transcriptome

Cells were grown in M9 medium with 11.3 mM NAG before being switched onto 22.5 mM malate with or without 3 mM of acetate. Samples were taken 30 min after the switch. Sample preparations and transcriptomics analyses were carried out as described by (Millard *et al*., 2021). Five independent replicates were analyzed for each condition. To analyze the transcriptomic data, a preliminary preprocessing step was performed using Excel to normalize the different conditions. The data were then processed through the EcoCyc platform, where Gene Ontology analysis was conducted to identify the most significant expressed and downregulated bioprocesses based on p-values. The transcriptomics data can be downloaded from the ArrayExpress database (www.ebi.ac.uk/biostudies/arrayexpress) under accession number E-MTAB-16650.

## Notes

https://www.ebi.ac.uk/biostudies/ArrayExpress/studies/E-MTAB-14542

https://www.ebi.ac.uk/biostudies/ArrayExpress/studies/E-MTAB-16650

### Competing Interest Statement

The authors have declared no competing interest.

### Summary of Updates

The revised manuscript broadens the evidence for acetate's beneficial effect by adding a glucose-to-fumarate transition tested in four E. coli strains, including Nissle 1917. New experiments showing that fumarate, succinate and malate also shorten lag, together with transcriptomic analysis of the Δacs mutant, strengthen the link to acetate assimilation and central metabolism. The Introduction and Discussion have been refocused, experimental choices clarified, and mechanistic claims qualified. Figures and legends have also been revised to improve readability and data presentation.

