## Supplementary figures for "Acetate promotes nutritional adaptation in *Escherichia coli*"

### **Figure S1 :** glucose to glucose adaptation control

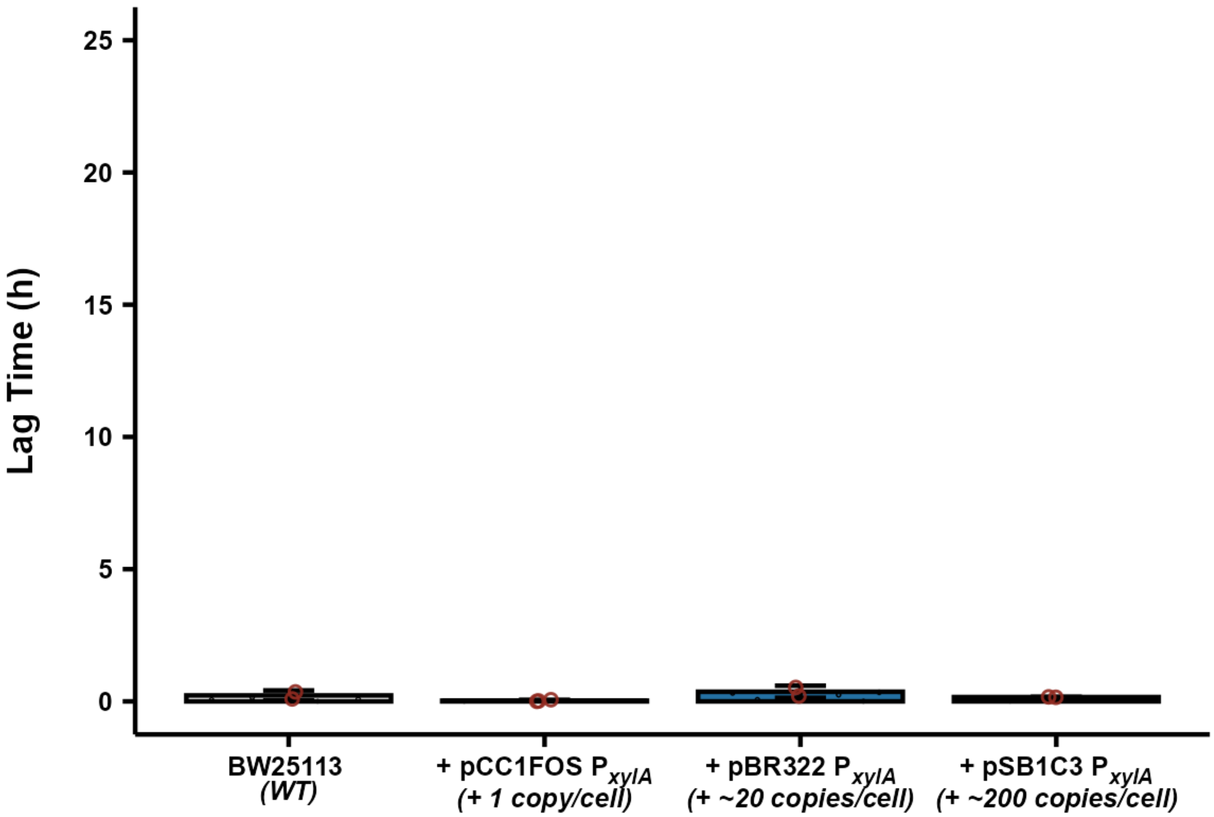


**Figure S1: lag times after a switch from to glucose to glucose of *E. coli* strains with different degrees of impaired xylose adaptation (see legend of Figure 1).** Each individual data point is represented by a red circle. The Y axis is the same as in Figure 1. Error bars represent standard deviations (SD).

- No significant lag was observed, proving that the lags observed in Figure 1 during the adaptation from glucose to xylose are purely physiological and not technical.

### **Figure S2:** most highly expressed genes in the glucose + xylose condition (M) compared with glucose alone (G).

| Syst. Name | Com. Name | M/G | Pvalue | X/G | Pvalue |
|---|---|---|---|---|---|
| b4677 | yobI | 6.00 | 1.E+00 | 3.50 | 1.E+00 |
| b4671 | ykgR | 5.00 | 1.E+00 | 4.00 | 1.E+00 |
| b0989 | cspH | 4.81 | 4.E-01 | 0.73 | 1.E+00 |
| b1729 | ydjN | 3.46 | 8.E-01 | 1.00 | 1.E+00 |
| b2424 | cysU | 3.35 | 1.E+00 | 1.85 | 8.E-01 |
| b2425 | cysP | 3.28 | 1.E+00 | 1.99 | 8.E-01 |
| b3704 | rnpA | 3.19 | 1.E+00 | 0.67 | 7.E-01 |
| b4098 | phnJ | 3.10 | 1.E+00 | 7.80 | 8.E-04 |
| b3088 | alx | 2.95 | 1.E+00 | 0.94 | 1.E+00 |
| b1565 | ydfV | 2.78 | 1.E+00 | 1.63 | 1.E+00 |
| b2423 | cysW | 2.76 | 1.E+00 | 1.96 | 5.E-01 |
| b0365 | tauA | 2.68 | 5.E-01 | 3.80 | 4.E-03 |
| b0113 | pdhR | 2.65 | 1.E+00 | 0.73 | 7.E-01 |
| b4341 | yjiS | 2.56 | 1.E+00 | 1.38 | 1.E+00 |
| b3175 | secG | 2.38 | 1.E+00 | 1.05 | 1.E+00 |
| b3471 | yhhQ | 2.33 | 1.E+00 | 1.10 | 1.E+00 |
| b2422 | cysA | 2.31 | 8.E-01 | 1.71 | 4.E-01 |
| b1821 | mntP | 2.27 | 1.E+00 | 1.24 | 9.E-01 |
| b2912 | fau | 2.27 | 1.E+00 | 1.17 | 1.E+00 |
| b0874 | ybjE | 2.25 | 1.E+00 | 0.66 | 9.E-01 |
| b2886 | ygfS | 2.22 | 1.E+00 | 1.70 | 7.E-01 |
| b0398 | sbcD | 2.18 | 1.E+00 | 0.99 | 1.E+00 |
| b3452 | ugpA | 2.17 | 1.E+00 | 0.89 | 1.E+00 |
| b3565 | xylA | 2.15 | 4.E-01 | 437.70 | 3.E-100 |
| b3568 | xylH | 2.14 | 1.E+00 | 157.87 | 7.E-79 |
| b4177 | purA | 2.12 | 1.E+00 | 1.31 | 8.E-01 |
| b0957 | ompA | 2.11 | 1.E+00 | 1.65 | 6.E-01 |
| b0413 | nrdR | 2.11 | 1.E+00 | 1.24 | 9.E-01 |
| b4062 | soxS | 2.09 | 1.E+00 | 1.06 | 1.E+00 |
| b1689 | ydiL | 2.08 | 1.E+00 | 2.58 | 8.E-01 |
| b4345 | mcrC | 2.07 | 1.E+00 | 1.48 | 9.E-01 |
| b4115 | adiC | 2.07 | 1.E+00 | 1.97 | 6.E-01 |
| b4423 | ldrC | 2.06 | 1.E+00 | 2.59 | 4.E-01 |
| b0446 | cof | 2.06 | 1.E+00 | 1.02 | 1.E+00 |
| b2583 | yfiP | 2.06 | 1.E+00 | 0.83 | 1.E+00 |
| b1433 | ydcO | 2.05 | 1.E+00 | 1.30 | 9.E-01 |
| b0468 | ybaN | 2.04 | 1.E+00 | 1.01 | 1.E+00 |
| b4246 | pyrL | 2.04 | 1.E+00 | 1.52 | 8.E-01 |
| b4557 | yidD | 2.02 | 1.E+00 | 0.73 | 8.E-01 |
| b3633 | waaA | 2.02 | 1.E+00 | 0.71 | 8.E-01 |
| b1627 | rsxA | 2.02 | 1.E+00 | 1.04 | 1.E+00 |
| b2764 | cysJ | 2.01 | 1.E+00 | 1.28 | 9.E-01 |
| b0813 | rhtA | 2.01 | 1.E+00 | 1.42 | 8.E-01 |
| b3566 | xylF | 2.00 | 1.E+00 | 816.12 | 6.E-13 |
| b3707 | tnaC | 2.00 | 1.E+00 | 5.63 | 3.E-01 |
| b2003 | yeeT | 2.00 | 1.E+00 | 2.89 | 5.E-01 |

**Figure S2: list of the most highly more expressed genes in the glucose + xylose condition (M) compared to glucose alone (G)** The table lists these genes, their relative expression level and the associated p values. Data are also provided for the xylose (X) versus glucose (G) condition.

- The presence of xylose may have a weak, but non-significant, effect on gene expression during growth on glucose.

### **Figure S3:** effect of xylose presence on single cells

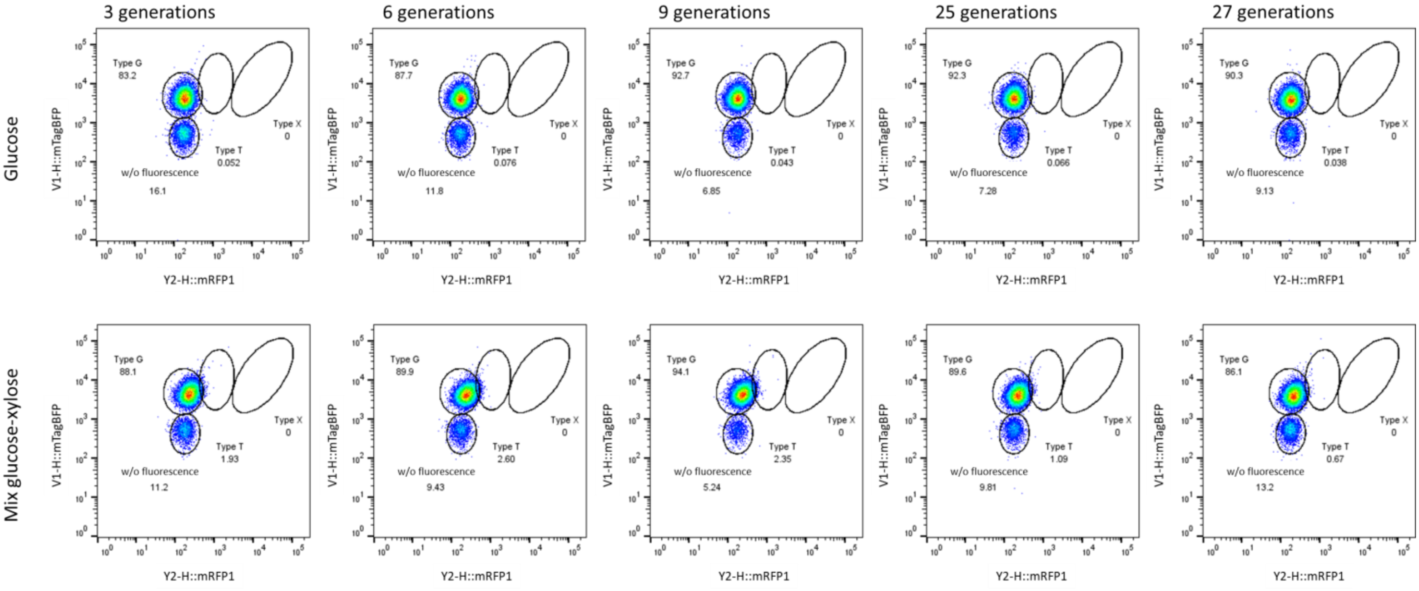


**Figure S3: cytometric profile of BW25113 pBR322-P$_{xylA}$-mRFP1 P$_{ihfB}$-mTagBFP cells maintained in exponential phasis for two days on glucose (top line) and glucose + xylose (bottom line).**
Cell distribution is represented in two dimensions (x-axis: red fluorescence of inducible mRFP1; y-axis: blue fluorescence of constitutive mTagBFP). The upper left lasso encompasses “glucose signature” cells, the upper right lasso “xylose signature” cells, and the middle lasso transitioning cells. The lower lasso corresponds to artifacts and cells that have potentially lost the plasmid.

- No significant increase in xylose-adapted cells was observed when xylose was present, even after 27 generations.

### **Figure S4 :** Effect of acetate during flask-based metabolic switches

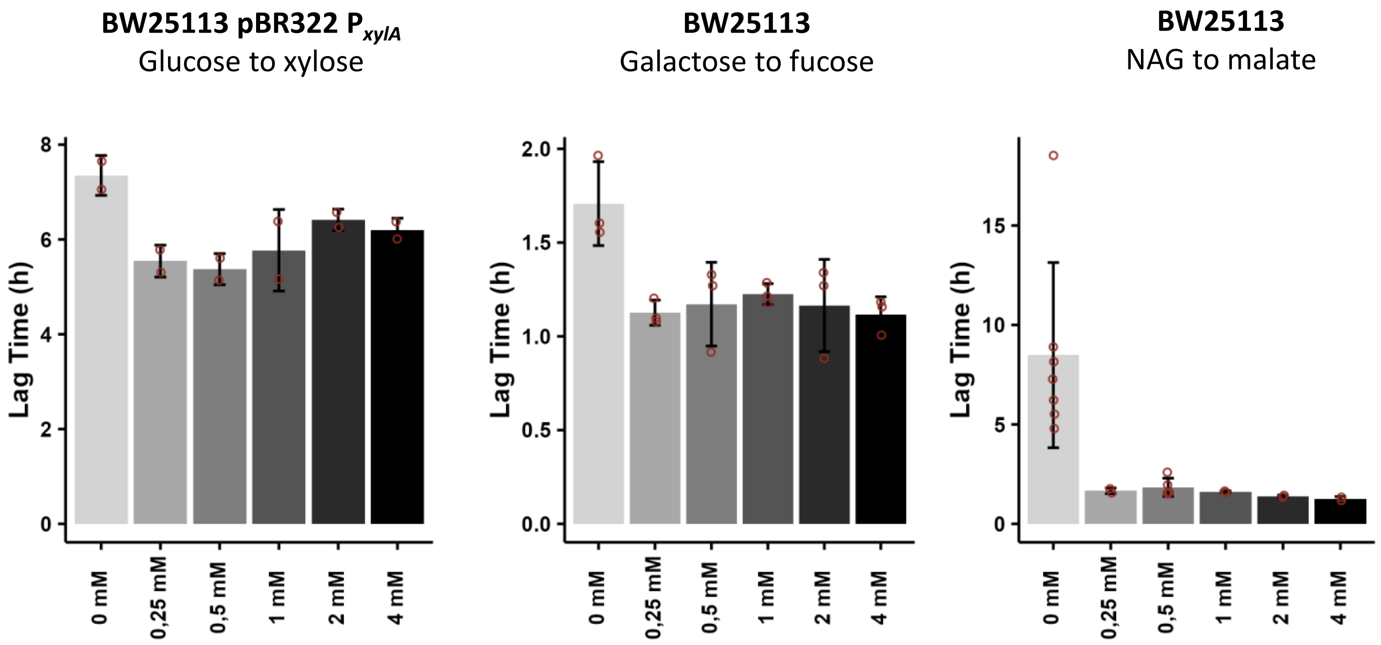


**Figure S4: acetate promotes metabolic adaptation in Erlen-Meyer flask.** Strain BW25113 carrying the titration plasmid pBR322 $P_{xylA}$ was switched from glucose to xylose (left graph). Wild type BW25113 was switched from galactose to fucose (mid graph) or from N-acetylglucosamine to malate (right part). 0 to 4 mM acetate was added in the second medium before adding the cells. All the cultures were performed in flask (2 to 3 repetitions). Each individual data point is represented by a red circle. Error bars represent standard deviations (SD).

- The same conclusion about the positive role of acetate could be drawn from the flask experiments as for the microplate assays presented in Figure 3.

**Figure S5 :** replicates of the galactose-fucose switches from Figure 4.

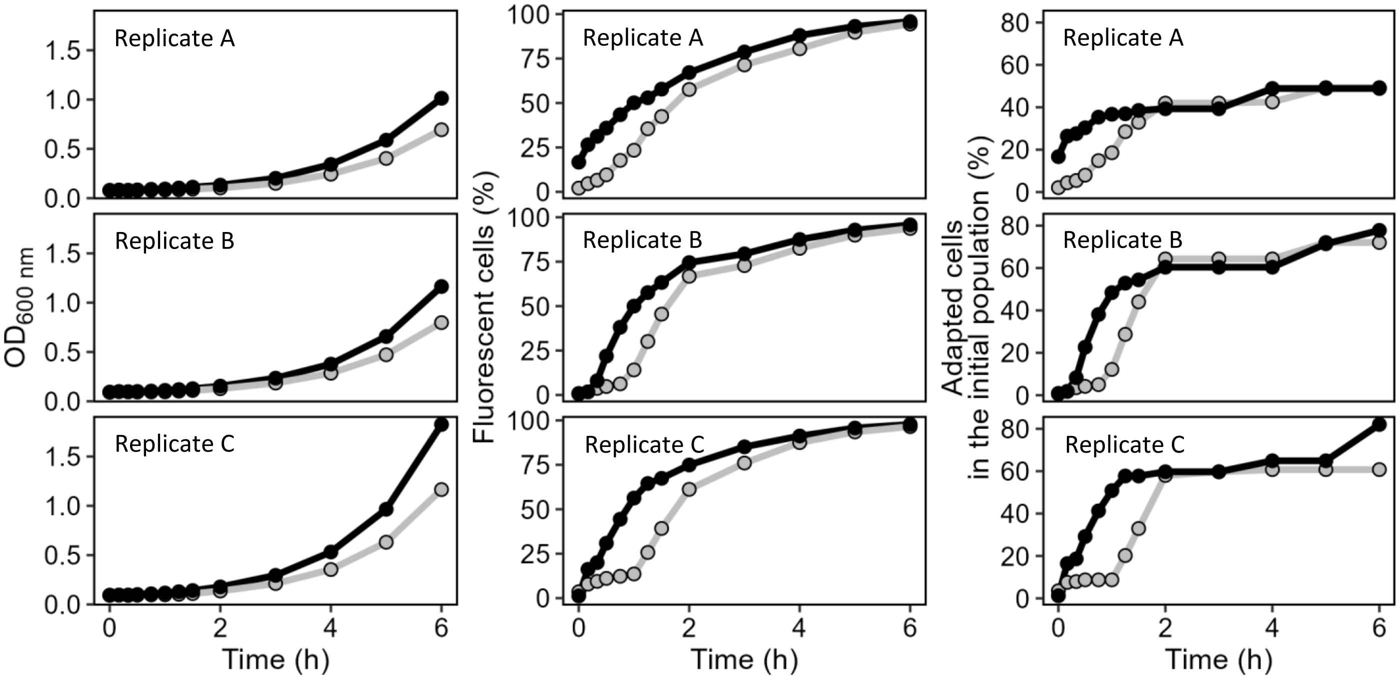


**Figure S5: replicates of the dynamics of total biomass** (left panel), **fluorescent subpopulations** (middle panel), and **percentages of adapted** cells in the initial population (right panel) during the switch of strain BW25113 pCC1FOS $P_{fucA}$-mNeonGreen $P_{j23119}$-mScarlet-I from galactose to fucose with or without 0.5 mM acetate (black and grey respectively)

### **Figure S6:** acetate consumption during switching from NAG to malate with an initial $OD_{600nm}$ = 0.1.

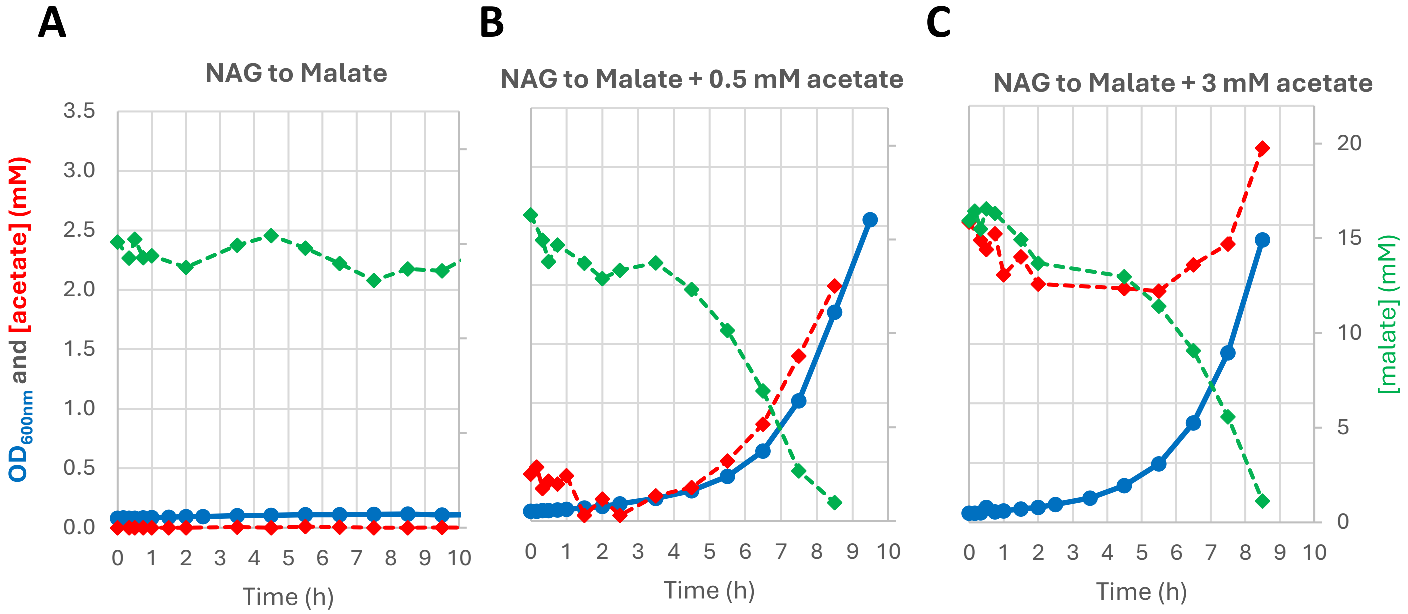


**Figure S6: acetate consumption during switching from NAG to malate with an initial $OD_{600nm}$ = 0.1.** Extracellular metabolites during switching from N-acetylglucosamine to malate in flasks. Biomass (optical density; blue dots), acetate concentration (red dots) and malate concentration (green dots) as a function of time, in the absence of acetate (A), with 0.5 mM acetate added to the malate (**B**) and with 3 mM acetate added to the malate (**C**).

- As observed under the 5B and 5C conditions, acetate is consumed by the cells well before any apparent resumption of growth on malate. The consumption rate is lower here, as this experiment was performed at an $OD_{600nm}$ of 0.1 rather than 0.6, as in Figure 5.

### **Figure S7:** adenylate charge with or without acetate

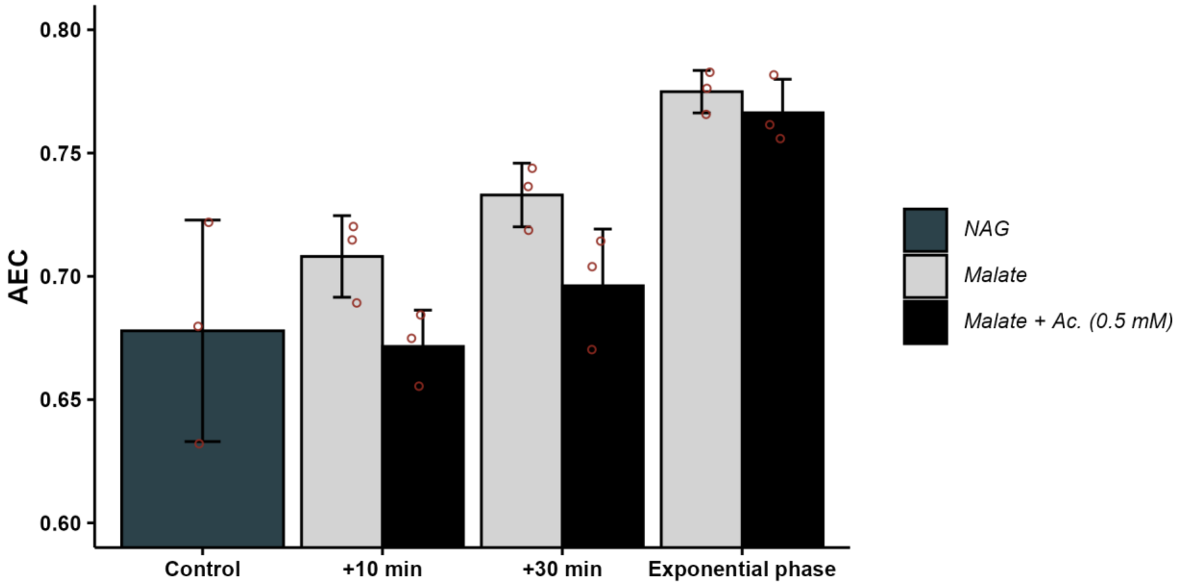


**Figure S7: comparison of adenylate energy charge during malate adaptation with or without acetate.** The same conditions and data as in Figure 5D were used here. Adenylate charge was calculated using the following formula:

$$AEC = \frac{[ADP] + \frac{1}{2}[ADP]}{[ATP] + [ADP] + [AMP]}$$

Control: exponential growth on N-acetylglucosamine, +10 minutes : 10 minutes after switching from N-acetylglucosamine to malate. +30 minutes: 30 minutes after switching from N-acetylglucosamine to malate. Exponential phase: during exponential phase on malate after switching from N-acetylglucosamine. These are the results of three independent experiments. Error bars represent standard deviations (SD).

- Acetate was suspected to increase the adenylate charge values, used here as an indicator of energy availability in the cell. However, the addition of acetate did not improve these values.

### **Figure S8 :** Effect of succinate, fumarate and malate during galactose to fucose switches

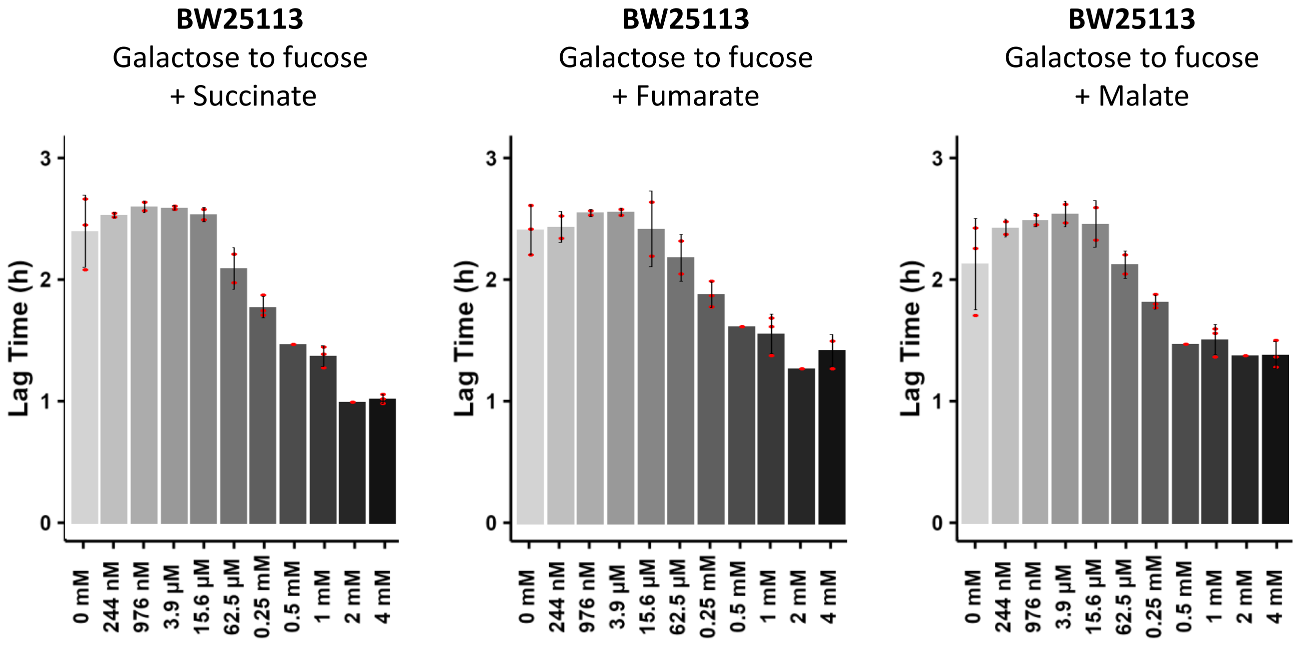


**Figure S8: other TCA cycle metabolites can promote nutritional adaptation.** Lag times were measured in *E. coli* BW25113 cultures switched from galactose to fucose in the presence of 0 to 4 mM succinate (left panel), fumarate (middle panel), or malate (right panel). N = 2 to 3 repetitions are shown as open red circles for each condition. Error bars represent standard deviations (SD).

- Although they are found in the extracellular medium at concentrations several orders of magnitude lower than those of acetate, other metabolites can promote the same effect when added at concentrations similar to that of acetate.

### **Figure S9:** no biological function are triggered by acetate in a *Δacs* mutant.

| **Ratio** | **Up-regulated** functions Cut Off = 3 | **p-value** | **Down-regulated** functions Cut Off = 0,33 | **p-value** |
|---|---|---|---|---|
| WT M+A / WT M | Biosynthetic process | 5.90e-9 | Biofilm formation * | 6.64e-5 |
| | Hydrogen sulphide biosynthetic process | 1.56e-6 | Hydrogen metabolic process | 8.56e-5 |
| | Nucleotide biosynthetic process * | 5.72e-10 | Methionine biosynthetic process * | 3.29e-10 |
| | Translation | 4.82e-7 | Response to stress * | 1.01e-8 |
| *ΔACS* M+A / WT M+A | Biofilm formation * | 2.10e-8 | IMP biosynthetic process | 4.19e-6 |
| | Methionine biosynthetic process * | 8.36e-5 | Nucleoside monophosphate metabolic process | 7.15e-7 |
| | Negative regulation of RNA biosynthetic process | 9.26e-6 | Nucleotide biosynthetic process * | 1.93e-5 |
| | Regulation of RNA biosynthetic process | 2.84e-6 | Ribonucleotide biosynthetic process | 9.66e-6 |
| | Response to stress * | 4.59e-7 | | |
| | Signaling | 5.00e-6 | | |
| *ΔACS* M+A / *ΔACS* M | No function found | / | No function found | / |
| *ΔACS* M / WT M | No function found | / | No function found | / |

**Figure S9: Gene ontology analysis of biological functions significantly enriched during growth resumption on malate.** A transcriptomic experiment was conducted to investigate gene expression 30 minutes after the switch from NAG to malate in either the wild-type BW25113 strain or its isogenic Δ*acs* mutant, in the presence or absence of 3 mM acetate (initial $OD_{600nm}$ = 0.6). Gene ontology analysis was then performed to identify cellular biological processes that were significantly up- or down-regulated.

- This highlights the strong response of the wild-type strain to acetate and its beneficial effect on growth resumption (Ratio WT M+A/WT M), as previously shown in Figure 3C.
- However, in the absence of acetate metabolism due to *acs* deletion, no cellular function was significantly enriched (Ratio Δ*acs* M+A/Δ*acs* M).
- We conclude that there appears to be no additional role for acetate beyond its metabolism that could explain its promoting effect on nutritional adaptation.
